# The CNS Microenvironment Promotes Leukemia Cell Survival by Disrupting Tumor Suppression and Cell Cycle Regulation in Pediatric T-cell Acute Lymphoblastic Leukemia

**DOI:** 10.1101/2023.09.01.555887

**Authors:** Sabina Enlund, Indranil Sinha, Christina Neofytou, Amanda Ramilo Amor, Konstantinos Papadakis, Anna Nilsson, Qingfei Jiang, Ola Hermanson, Frida Holm

## Abstract

A major obstacle in improving survival in pediatric T-cell acute lymphoblastic leukemia is understanding how to predict and treat leukemia relapse in the CNS. Leukemia cells are capable of infiltrating and residing within the CNS, where they interact with the microenvironment and remain sheltered from systemic treatment. These cells can survive in the CNS niche, by hijacking the microenvironment and disrupting normal functions, thus promoting malignant transformation. While the protective effects of the bone marrow niche have been widely studied, the mechanisms behind leukemia infiltration into the CNS and the role of the CNS niche in leukemia cell survival remain unknown.

We have identified a dysregulated gene expression profile in CNS infiltrated T-ALL and CNS relapse, promoting cell survival, chemoresistance and disease progression. Furthermore, we discovered that interactions between leukemia cells and CNS microenvironment induce epigenetic alterations, such as changes in gene regulation and histone modifications, including H3K36me3 levels.

These findings can be utilized to predict CNS infiltration and CNS relapse, therefore avoiding overtreatment and adverse effects caused by CNS directed therapy. Additionally, the identified genetic drivers of disease progression can serve as a first step towards identifying therapeutic targets, to sensitize the CNS niche to current therapeutic strategies.

## 1. INTRODUCTION

T-Cell acute lymphoblastic leukemia (T-ALL) is an aggressive hematological malignancy, caused by uncontrolled proliferation of T-cell progenitors. Although modern treatment has improved the overall survival, more than 20% of all children with T-ALL will relapse typically within 2 years of diagnosis [1]. Approximately 30% of all relapse cases occur in the central nervous system (CNS) and CNS involvement is a risk factor for treatment failure and increased mortality [2]. Clinically, CNS involvement in T-ALL is diagnosed by quantifying the number of white blood cells and the presence of lymphoblasts in the cerebrospinal fluid (CSF) ranging from CNS1 (no CNS infiltration) to CNS3 (advanced CNS infiltration). Despite the clear impact of CNS disease on survival of children with CNS involved T-ALL, the infiltration route, and underlying mechanisms of leukemia cell survival in the CNS niche is not fully understood [3].

Most children who relapse in the CNS are initially diagnosed as CNS1. In the light of this, new diagnostic measures are required to properly diagnose and identify T-ALL patients at risk of CNS relapse. To date, CNS directed therapy is given as a prophylactic treatment to prevent CNS relapse, regardless of CNS involvement at the time of diagnosis. However, CNS directed therapy causes detrimental adverse effects and neurocognitive impairments, as well as increasing the risk of secondary malignancies in the CNS [4]. Due to lack of knowledge of the mechanisms promoting leukemia survival in the CNS microenvironment, no new CNS directed treatments have been developed since the 1960’s, yet the treatment is considered essential for complete cure [4]. Thus, there is a need for reliable biomarkers to properly predict and monitor disease progression and CNS involvement to identify high risk patients, thus minimizing the risk of overtreatment and neurotoxic effects.

Through RNA-sequencing analysis of pediatric T-ALL patients samples (pre-generated by the TARGET initiative), both at time of diagnosis and following relapse in the CNS, we have identified a differential gene expression pattern during different stages of CNS involvement. Therefore, we hypothesize that CNS disease in primary T-ALL and CNS relapse are typified by distinct genetic changes which could be utilized as predictive markers to properly diagnose patients at risk of relapsing in the CNS. We discovered a potential role of histone lysine methyltransferase SETD2 in regulating tumor suppression in CNS relapsed T-ALL patients, which was further endorsed by SETD2 manipulation in T-ALL cells, causing dysregulation of the TP53 pathway as well as several genes involved in self-renewal and survival. Furthermore, we report that genetic alterations caused by interactions between leukemia cells and the CNS microenvironment promote leukemia cell survival and chemoresistance, thus promoting disease progression. To test this hypothesis, T-ALL cells were cultured together with a glioma cell line, to identify genetic and epigenetic changes caused by direct interactions of leukemia cells and CNS derived cells. Our analysis revealed dysregulation of several pathways capable of promoting malignant progression as well as disruption of tumor suppression via p53 signaling. T-ALL cells cultured adherent to glioma cells displayed loss of TP53 expression and an upregulation of several pro-survival genes, including the long, anti-apoptotic BCLX isoform. To further study the effect of CNS co-culture on T-ALL survival, cells were treated with Methotrexate (MTX), a commonly used drug with excellent CNS penetration characteristics, in co-culture and in suspension followed by RT-qPCR analysis. We found dysregulation of several cell cycle regulatory genes, potentially inducing senescence during chemotherapeutic treatment, thus promoting therapy resistance.

Taken together, our findings indicate a differential gene expression pattern in CNS involved T-ALL, which could potentially be utilized as predictive biomarkers to properly identify patients at risk of CNS relapse. Our model of the CNS niche induced alterations in leukemia cells promoting survival and chemoresistance, possibly serving as future therapeutic targets.

## 2. METHODS

### 2.1. RNA sequencing analysis

30 diagnosis T-ALL and 18 Relapse T-ALL samples were selected for this analysis, samples are not longitudinal. The RNA sequencing dataset (.BAM) files sourced from T-ALL patients were acquired via the GDC Data Portal (https://portal.gdc.cancer.gov/projects/TARGET-ALL-P2). The gene expression profiles were established using the SeqMonk Mapped Sequence Data Analyzer tool (version 1.47.1), resulting in Frames per Kilo-base of exon per Million mapped reads (FPKM) values. The RNA-seq quantitation pipeline was used to the subsequent criteria: Transcript features were set to mRNA, Library type was selected as Opposing strand specific and Transcript isoforms were merged. Normalized values equal to or less than 0.1 (<=0.1) were transformed into “NA,” while the remaining values were converted to the log2 scale. The resultant file was employed for subsequent analyses.

To perform Gene-Set Enrichment Analysis (GSEA), we used data generated through RNA-seq quantitation pipeline. This analysis utilized GSEA version 4.2.3 (build 10) along with the KEGG gene set database (c2.cp.kegg.v2023.1.Hs.symbols.gmt).

To analyze gene isoforms, sample sequences were downloaded from GDC data portal. GenomicDataComons package and a R script was used (R version 4.0.3). SeqMonk was used to generate FPKM values.

### 2.2. Cell lines

T-ALL cell lines CCL-119, Jurkat and SUP-T1 (ATCC) were cultured in suspension in RPMI with 10% FBS (Sigma-Aldrich). T-ALL cell line MOLT16 (Leibniz Institute DSMZ) was cultured in suspension IMDM (Gibco) with 10% FBS. Glioblastoma cell line U87-MG (ATCC) was maintained as adherent culture in ATCC modified Eagle’s Minimum Essential Medium (ATCC) passaged by detachment using 0.25% trypsin-EDTA (Gibco). Media renewal and passaging was performed every 2-3 days and cells were incubated at 37 °C with 5% CO2.

### 2.3. Co-culture

U87-MG cells were plated and allowed to adhere in EMEM, reaching 60-70% confluency. T-ALL cells were added to the wells at ratio of 3:1 leukemia cells to glioblastoma cells. Non-adherent cells were collected after 24 hours followed by media renewal. MTX (Sigma Aldrich) was resuspended and diluted in DMSO (Sigma) and added to the cells in suspension and co-culture 24 hours after plating at a concentration of 2μM. After an additional 48 hours non-adhered cells were discarded and adhered T-ALL cells were separated from glioblastoma cells using EasySep Human CD3 positive Selection kit II (Stem Cell Technologies) and viability was determined by trypan blue staining (Bio-Rad). Control conditions included T-ALL cells in RPMI or IMDM, T-ALL cells in EMEM and U87-MG in EMEM separately.

### 2.4. RNA preparation and qRT-PCR analysis

Total RNA was extracted from cells with RNeasy Micro kit (Qiagen) according to the provided protocol. First-strand complementary DNA (cDNA) synthesis was performed using the Invitrogen SuperScript III First-Strand synthesis SuperMix for qRT-PCR (Invitrogen), using 500 ng template RNA. Quantitative real-time polymerase chain reaction analysis (qRT-PCR) was performed using SYBR GreenER qPCR SuperMix for iCycler (Invitrogen) following suppliers recommended program. Target messenger RNA (mRNA) was normalized to hypoxanthine phosphoribosyl transferase (HPRT) mRNA transcript levels and fold change were calculated via the delta-delta cycle threshold (CT) method normalizing mean delta CT of each cell line to the delta CT of their respective control. Primer sequences can be found in supplementary table S1.

### 2.5. Chromatin Immunoprecipitation

ChIP was performed using the Diagenode True Microchip kit (Diagenode), following the manufacturer’s protocol. Cells were collected at 100.000 cells per immunoprecipitation, including histone 3 (H3) as a positive control, immunoglobulin G (IgG) as a negative control, histone mark trimethylation of lysine 36 on histone 3 (H3K36me3) and histone mark trimethylation of lysine 27 on histone 3 (H3K27me3). ChIP-qPCR was performed using SYBR GreenER qPCR SuperMix for iCycler (Invitrogen) according to the supplier’s program. Data is presented as % Input (100*2ΔCT).

### 2.5. Plasmid cloning

DH5α competent cells (Invitrogen) were transformed by heat shock in 42°C water bath using SETD2 Human Tagged ORF Clone (Origene, RG224760) or pCMV6□AC□GFP Mammalian Expression Vector (Origene, PS100010) followed by overnight culture on LB agar plates (Sigma-Aldrich) containing 100μg/ml ampicillin (Sigma-Aldrich) at 37°C. Plasmid DNA cloning and purification was performed using QIAGEN Plasmid Midi Kit (Qiagen) according to manufacturer’s protocol.

### 2.6. Transfections

Cell□line□specific Nucleofector Kits (Lonza) were used for transfections which were performed according to manufacturer’s protocol on a Nucleofector® 2b Device (Lonza). Nucleofection efficiency was determined by RT□qPCR 24 hours post transfection.

### 2.7. Statistical analysis

Statistical analyses were performed using either unpaired t-test, two-way ANOVA or Pearson correlation coefficients, two-tailed with 95% confidence interval. Significance was set to P≤0.05 (*P ≤ 0.05, **p<0.01, ***p<.001, ****p<0.0001). Results are presented as the mean ± SEM. Outliers statistically excluded by Robust regression and Outlier removal (ROUT).

## 3. RESULTS

### 3.1. CNS infiltrated T-ALL and CNS relapsed T-ALL patients are distinguished by differential gene expression patterns compared to patients at time of diagnosis without CNS involvement

To determine alterations of gene expression in pediatric T-ALL patients with CNS involvement and patients who have relapsed in the CNS, a total of 48 samples were included, from patients at time of diagnosis and patients with relapsed T-ALL. Inclusion criteria was based on number of white blood cells (WBCs), MLL status (negative), age and tissue of origin (bone marrow) (Figure 1A). Samples taken from patients at initial diagnosis were subdivided depending on the presence of blasts and number of WBCs in the CSF, ranging from CNS1 (no CNS infiltration) to CNS3 (advanced CNS infiltration) and relapsed patients were divided by site of relapse (CNS or other) (Figure 1B). Whole genome RNA-sequencing analysis revealed differences between all subgroups of patients, with some genes commonly dysregulated through several groups with CNS disease (Figure SF1A). Samples from patients relapsed in the CNS were distinguished from patients at time of diagnosis, both classified as CNS1 and CNS3 (Figure 1C). In CNS3 patients with primary T-ALL and patients who relapsed in the CNS, there were less than 2% genes significantly expressed compared to patients diagnosed as CNS1 at time of diagnosis (Figure 1D, Figure SF1B). However, in patients with CNS relapse most genes were significantly upregulated compared to CNS1 diagnosis patients, while in diagnosis patients classified as CNS3, genes were generally downregulated (Figure 1D, Figure SF1B). Furthermore, both patients diagnosed as CNS3 at time of diagnosis and CNS relapse patients displayed a distinct gene signature compared to CNS1 diagnosis patients (Figure 1E, SF1C). Interestingly, samples from CNS relapsed patients revealed an upregulation of cell cycle inhibitor CDKN2A [5] and tumor suppressor RASSF8 [6] (Figure 1F, Figure SF1D). Several genes from the HOX family of transcription factors were upregulated in CNS relapse, including HOXA3 and HOXB2, which have been shown to support immortalization of leukemia cells in acute myeloid leukemia (AML) (Figure 1F) [7].

**Figure 1.**
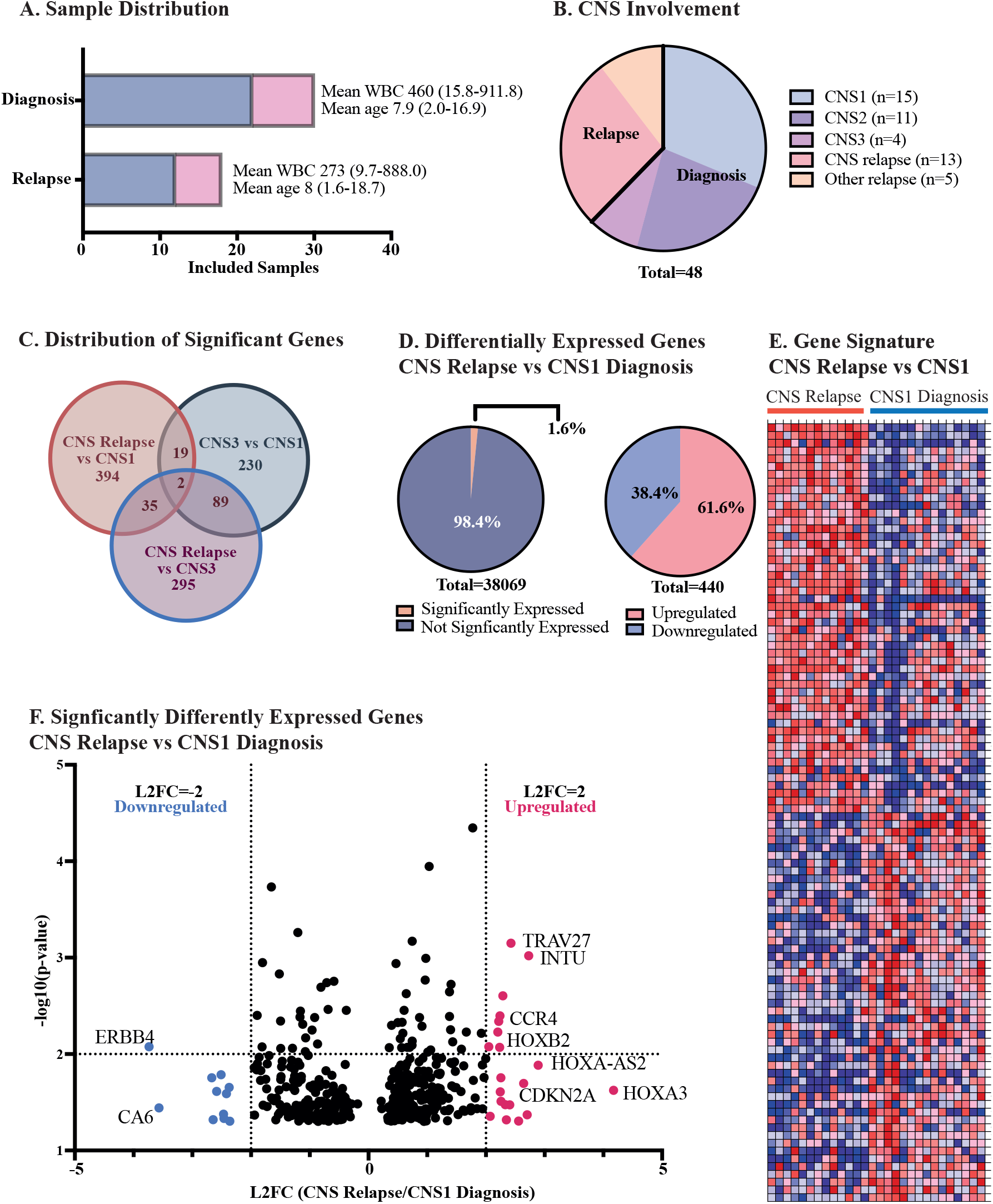
Gene signature of CNS relapsed T-ALL. **A)** Schematic overview of included T-ALL patients from the TARGET database. Samples were distributed from time of diagnosis (n=30) and relapse (n=18) and include only bone marrow samples. Ratio of male:female patients is 22:8 for diagnosis patients with a mean WBC of 460 and mean age of 7.9. Relapsed patients had male:female ratio of 12:6 with a mean WBC of 273 and mean age of 8. **B)** Degree of CNS involvement of diagnosis samples based on number of WBCs and presence of blasts in the CSF ranging from CNS1 (no CNS disease, n=15) to CNS2 (mild CNS disease, n=11) to CNS3 (advanced CNS disease, n=4). Relapse patients are subdivided by site of relapse, CNS relapse (n=13) or other site of relapse (n=5). **C)** Venn diagram showing distribution of significantly expressed genes (p<0.05) in samples with CNS relapse versus CNS1 and CNS3 diagnosis samples (without novel transcripts). **D)** Percentage of significantly differentially expressed genes (left pie chart) and distribution of up and downregulated genes (right pie chart) in CNS relapse compared to diagnosis without CNS involvement. **E)** Heat map of 50 differentially expressed genes in CNS relapse versus CNS1 diagnosis samples, data displayed as log values collapsed to symbols. **F)** Volcano plot of significantly differentially expressed genes in CNS relapse versus CNS1 diagnosis samples, displayed as L2FC calculated from normalized FPKM values (p<0.05).

### 3.2. Disruption of tumor suppression via the TP53 pathway and apoptosis in CNS involved T-ALL and CNS relapsed T-ALL patients

To further elucidate the impact of CNS infiltration on T-ALL patients, GSEA analysis was performed to identify dysregulated pathways and gene sets in CNS relapse compared to patients diagnosed as CNS3 at initial diagnosis. Analysis revealed an enrichment of cell adhesion molecules and several key pathways in promoting self-renewal and hematopoietic stem cell maintenance (NOTCH and WNT signaling pathways) (Figure SF2A-C) [8]. CNS relapse upregulates cell adhesion molecule N-Cadherin, which was reported to promote therapy resistance of leukemia cells through microenvironmental interactions (Figure 2A) [9]. Moreover, expression levels of β-catenin were downregulated in CNS3 diagnosis patients and CNS relapse compared to CNS1 diagnosis patients (Figure 2A). β-catenin, as part of the canonical Wnt/β-catenin pathway, is involved in regulating cell growth and proliferation [10]. RNA-sequencing analysis also identified dysregulation of RASSF2, a K-Ras effector protein considered a tumor suppressor with pro-apoptotic capacities often inactivated in cancer [11]. Interestingly, RASSF2 was found to be downregulated in CNS3 diagnosis pagtients yet upregulated in CNS relapse, compared to CNS1 diagnosis patients (Figure 2A). GSEA also identified an enrichment of the P53 pathway in CNS relapse, responsible for tumor suppression by regulating apoptosis, DNA repair and senescence (Figure 2B) [12]. A downregulation of proto-oncogene MDM2 was observed in CNS infiltrated T-ALL samples, responsible for controlling TP53 activity (Figure 2A) [12]. Furthermore, CNS relapse patients were distinguished by a dysregulation of the apoptotic pathway, with a downregulation of the apoptotic regulator BCL2 (Figure SF2D-E). While no significant differences were found in expression levels of BCL2L1 (BCLX), differential expression of splice isoforms was prevalent, with an upregulation of the long, pro-survival isoforms of both BCL2 and BCLX (BCL2L and BCLxL) in almost all T-ALL patient samples, suggesting a pro-survival isoform switch during T-ALL progression at all stages of CNS involvement (Figure 2C, Figure SF2E) [13].

**Figure 2.**
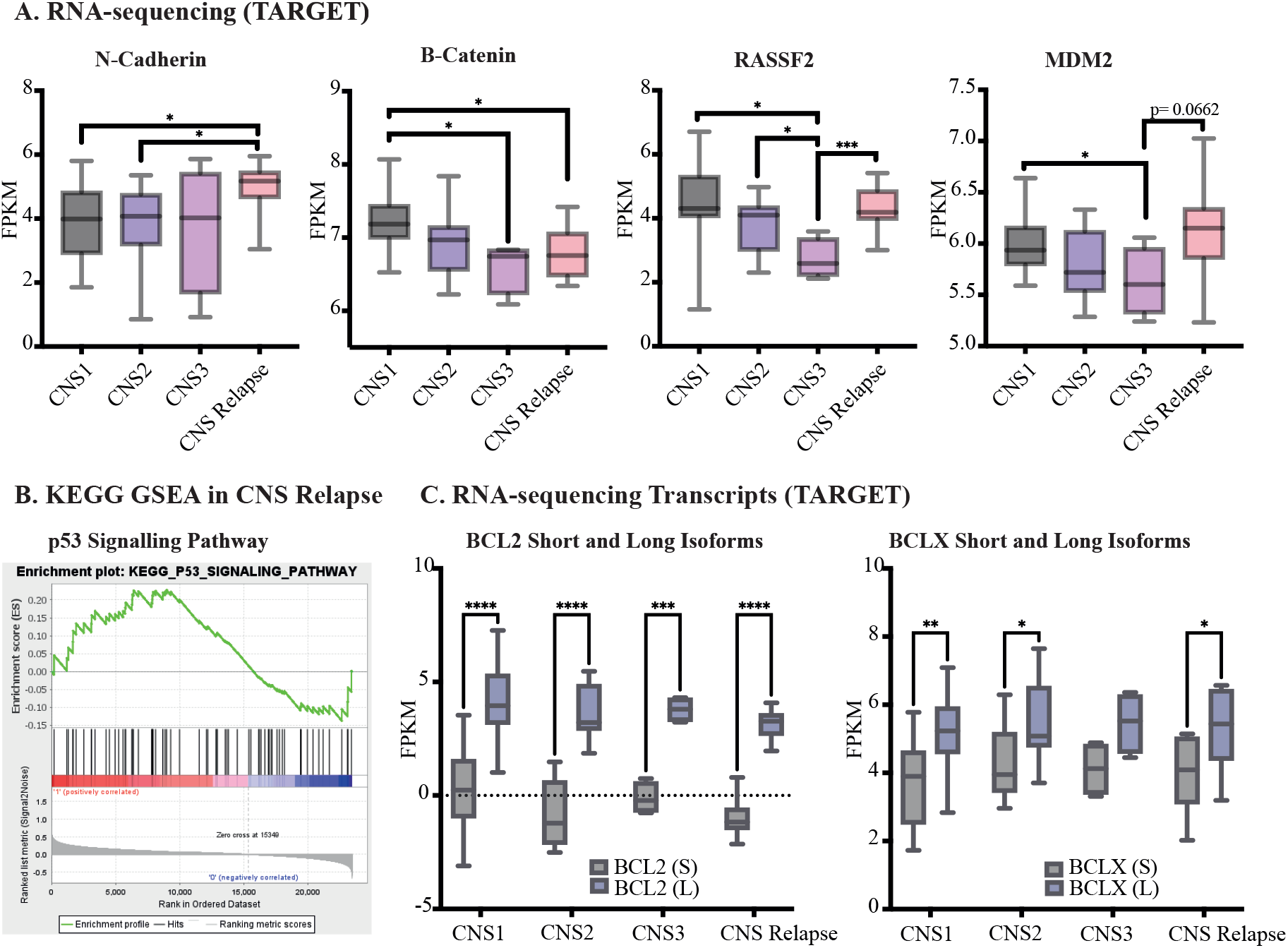
Enrichment plots and dysregulated genes in CNS involved T-ALL. Gene and isoform expression data is presented as normalized FPKM values obtained from the TARGET database from T-ALL patients at time of diagnosis divided by degree of CNS involvement and CNS relapsed T-ALL patients. **A)** RNA-sequencing analysis of normalized FPKM values showing differential gene expression of N-Cadherin, B-catenin, RASSF2 and MDM2 in CNS involved T-ALL and CNS relapsed T-ALL. (Significance calculated by two-tailed, unpaired Student’s t-test. *p<0.05, ***p<0.001). **B)** GSEA enrichment plot of p53 signalling pathway in CNS relapse versus CNS1 diagnosis patients, showing the profile running ES score and positions of gene set members on the ranked-ordered list. GSEA was performed using KEGG pathways. **C)** RNA-sequencing analysis of normalized FPKM values showing increased expression of BCL2 and BCLX long, pro-survival isoforms in T-ALL diagnosis samples and CNS relapsed T-ALL. (Significance was calculated by two-way ANOVA. *p<0.05, **p<0.01, ***p<0.001, ****p<0.0001).

### 3.3. SETD2 is involved in regulating tumor suppression and apoptosis in T-ALL

T-ALL patients with CNS disease were typified by a dysregulation of genes involved in epigenetic regulation of transcription and chromatin organization, including a downregulation of the lysine acetyltransferase KAT6A and histone methyltransferase SETD2 (Figure 3A, Figure SF3A). SETD2 is commonly mutated in cancer and is involved in regulating transcriptional activation, splicing and mismatch repair by trimethylating lysine 36 on histone 3 (H3K36me3) [14]. Although SETD2 has previously been recognized as a potential tumor suppressor by interacting with p53, the role of SETD2 and the effects of the histone mark H3K36me3 in leukemia is unclear [14, 15]. Interestingly, RNA-sequencing analysis revealed a negative correlation between SETD2 and TP53 in CNS relapse patients, which cannot be seen in CNS1 patients at diagnosis of T-ALL (Figure 3B, Figure SF3B). Furthermore, SETD2 and MDM2 were found to have a positive correlation in CNS relapse which could not be observed in CNS1 patients at primary diagnosis (Figure 3B, Figure SF3B). STRING analysis revealed a direct protein-protein interaction between SETD2 and TP53, as well as an indirect link between SETD2 and MDM2, potentially via EP300 and SIRT1 (Figure 3C). Both EP300 and SIRT1 were found to be downregulated in CNS involved T-ALL and with a positive correlation to SETD2 in all T-ALL samples (Figure SF3C-D) [16]. Additionally, STRING analysis identified direct and indirect interactions between SETD2 and NANOG, SOX2 and OCT4, known for regulating pluripotency and self-renewal [17, 18], as well as apoptotic regulators BCL2 and BCLX (Figure 3C).

**Figure 3.**
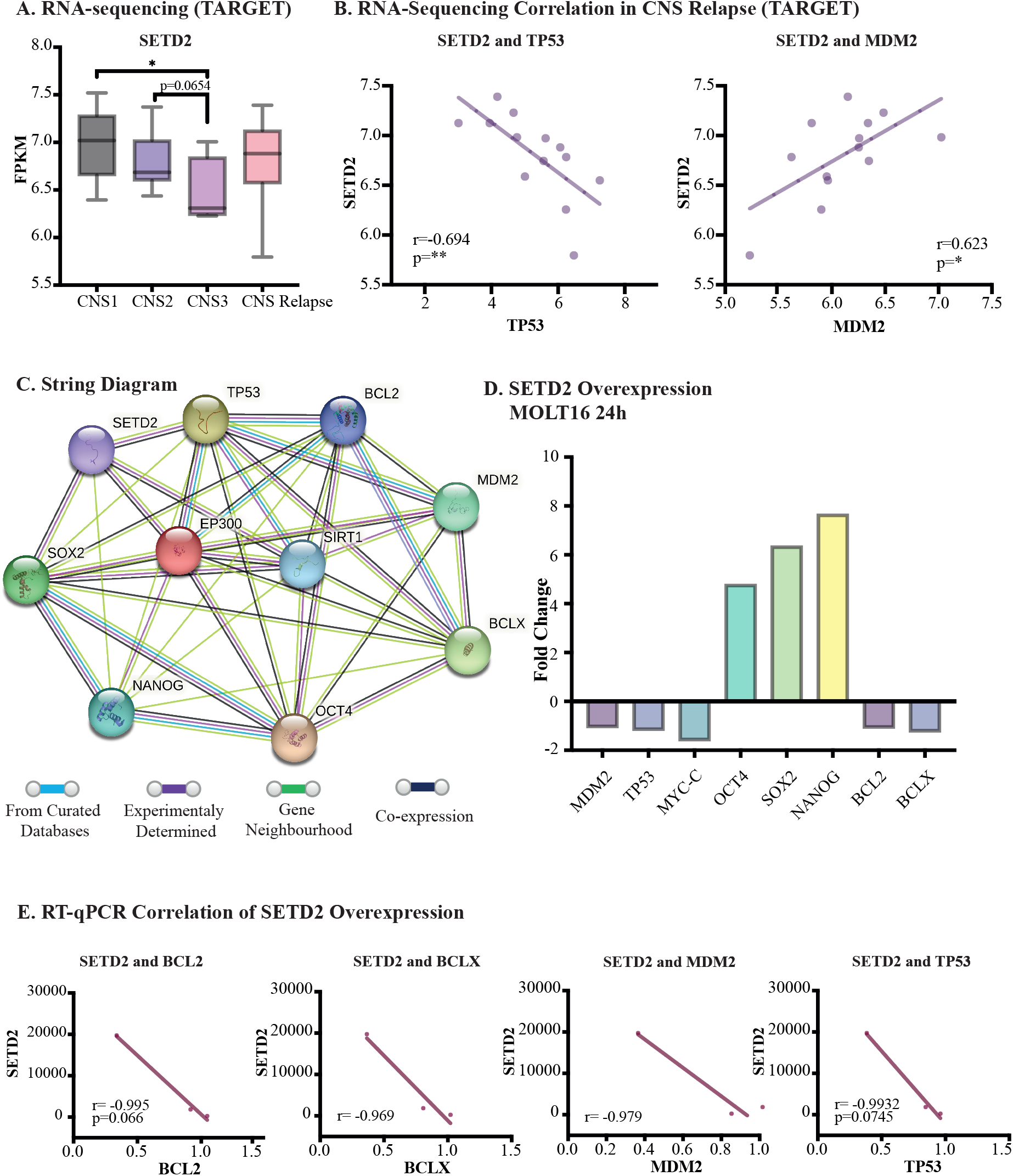
The role of SETD2 in tumor suppression and apoptosis. **A)** RNA-sequencing analysis of normalized FPKM values showing decreased expression of SETD2 in CNS3 diagnosis patients compared to CNS1 diagnosis samples. (Significance calculated by two-tailed, unpaired Student’s t-test. *p<0.05). **B)** RNA-sequencing correlation analysis of SETD2 and TP53 (left graph) showing a negative correlation (r=-0.694, **p=<0.01) and MDM2 (right graph) showing a positive correlation (r=0.628, *p=<0.05) in CNS relapsed T-ALL patients. (Correlation was calculated using Pearson correlation coefficients, two-tailed with 95% confidence interval). **C)** String diagram showing protein-protein interactions directly and indirectly experimentally linking SETD2 with TP53, MDM2, BCLX, BCL2, NANOG and OCT4. **D)** mRNA expression levels (fold change versus empty vector control) of MDM2, TP53, MYCC, OCT4, NANOG, BCL2 and BCLX 24 hours after SETD2 overexpression by transfection in MOLT16 cells. **E)** mRNA expression levels of SETD2 overexpression in T.-ALL cell lines MOLT16, Jurkat, CCL-119 negatively correlates with BCL2 (r=0.-995, p=0.066), BCLX (r=-0.969), MDM2 (r=-979) and TP53 (r=-0.993, p=0.075). (Correlation was calculated using Pearson correlation coefficients, two-tailed with 95% confidence interval).

To experimentally test the effects of SETD2 on tumor suppression and cell survival, T-ALL cell lines were transfected with a vector designed to overexpress SETD2 (Figure SF3E-F). Overexpression of SETD2 led to a dysregulation of several genes involved in tumor suppression, self-renewal, and apoptosis (Figure 3D). Interestingly, SETD2 overexpression caused a downregulation of TP53, MDM2, MYC-C, BCL2 and BCLX, while OCT4, NANOG and SOX2 were upregulated (Figure 3D). Moreover, SETD2 overexpression levels correlate negatively with both BCL2 and BCLX whole gene expression, as well as with MDM2 and TP53 (Figure 3E). SETD2 manipulation also affected expression levels of BCL2 and BCLX isoforms, causing decreased expression of the BCLX short, pro-apoptotic isoform and an increased expression of the long, pro-survival isoform (Figure SF3G). Together, this data suggests a regulatory role of SETD2 in survival and self-renewal of T-ALL cells; SETD2 is potentially involved in disrupting TP53 and MDM2 expression levels in CNS relapsed T-ALL patients.

### 3.4. CNS co-culture inhibits tumor suppression and promotes cell cycle arrest during methotrexate treatment

To further study the role of the CNS niche on leukemia cell survival and gene program regulation, a 72-hour co-culture system with T-ALL cell lines and a glioblastoma cell line was developed. CNS co-culture revealed dysregulation of several genes in T-ALL cells, including increased expression of cancer stem cell marker CD44 [19, 20] and the adhesion molecule ICAM1 [21], as well as β-catenin and NANOG, compared to cells grown alone in suspension (Figure 4A, Figure SF4A). Moreover, tumor suppressor TP53 was significantly downregulated in T-ALL cells grown in co-culture, while proto-oncogene MDM2 was upregulated compared to cells in suspension (Figure 4A). Although no significant difference could be observed in SETD2 mRNA levels, changes to the SETD2 generated histone mark H3K36me3 could be observed following co-culture (Figure 4B, Figure SF4A). Chromatin Immunoprecipitation coupled with RT-qPCR revealed an enrichment of H3K36me3 on the promoter region of MDM2 (Figure 4B). Furthermore, H3K36me3 was enriched on the promoter region of BCLX, correlating with increased mRNA expression levels of both BCLX whole gene and the BCLX long, pro-survival isoform in T-ALL cells grown in CNS co-culture compared to cells in suspension (Figure 4C, Figure SF4B-C).

**Figure 4.**
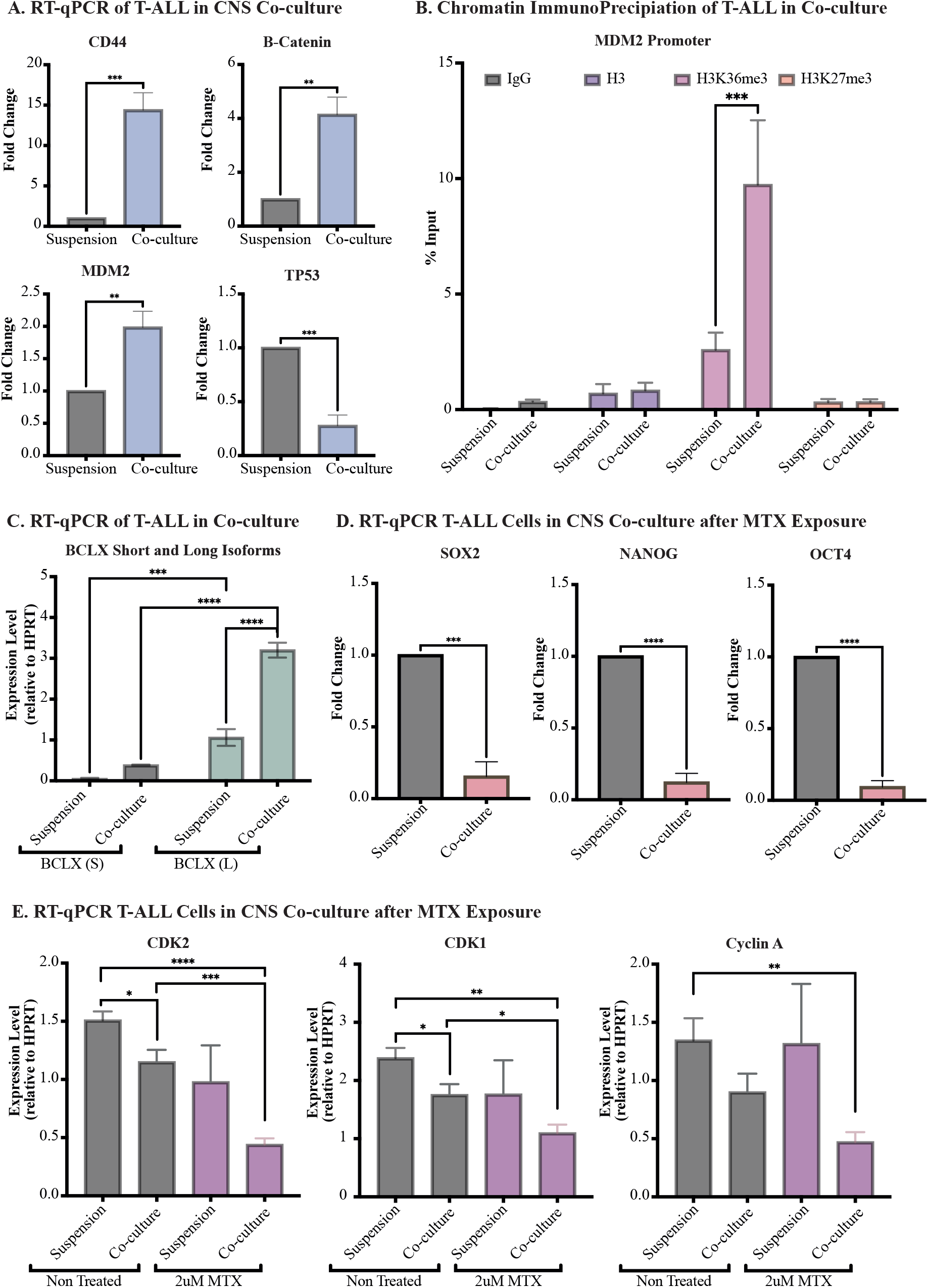
CNS co-culture disrupts tumor suppression and cell cycle regulation in T-ALL cells. T-ALL cell lines MOLT16, Jurkat, CCL-119 and SUP-T1 were cultured adherent to glioblastoma cell line U87-MG for 72 hours. **A)** RT-qPCR analysis of T-ALL cell lines in CNS co-culture showing increased expression (fold change) of CD44, B-catenin and MDM2, as well as downregulation of TP53 (outlier statistically excluded by ROUT) compared to cells grown in suspension. (Significance calculated by two-tailed, unpaired Student’s t-test. **p<0.01, ***p<0.001). **B)** Chromatin Immunoprecipitation coupled with RT-qPCR showing increased expression of H3K36me3 on the MDM2 promoter region in T-ALL cell lines grown in CNS co-culture compared to cells grown in suspension. (Significance was calculated by two-way ANOVA, ***p<0.001). **C)**RT-qPCR analysis of T-ALL cell lines in CNS co-culture showing increased expression (relative to HPRT) of BCLX long, pro-survival isoform compared to cells grown in suspension. (Significance was calculated by two-way ANOVA, ***p<0.001, ****p=<0.0001). **D)**RT-qPCR analysis of T-ALL cells in CNS co-culture after 48 hours of 2uM MTX treatment, showing decreased expression of SOX2, NANOG and OCT4 compared to T-ALL cell lines treated with 2uM MTX in suspension. MTX=methotrexate. (Significance calculated by two-tailed, unpaired Student’s t-test. ***p<0.001, ****p=<0.0001). **E)** RT-qPCR analysis of T-ALL cell lines in CNS co-culture and suspension with and without 48 hours of 2uM MTX treatment, showing decreased expression (relative to HPRT) of CDK2 and CDK1 in non-treated and MTX treated T-ALL cell lines in grown co-culture, as well as decreased expression of cyclin A2 in MTX treated T-ALL cell lines grown in co-culture compared to cells grown in suspension. (Significance calculated by two-tailed, unpaired Student’s t-test. *p<0.05, **p<0.01, ***p<0.001, ****p=<0.0001).

Previous research has shown that T-ALL cells grown in co-culture and in direct contact with CNS derived cells display increased survival and resistance to therapy compared to cells grown in suspension when treated with chemotherapy [22, 23]. However, mechanisms promoting increased survival are not well understood. Therefore, we exposed T-ALL cells to MTX, a chemotherapeutic agent commonly used to treat CNS disease in T-ALL [4]. Cells were treated with 2μM of MTX, both in suspension and when grown adherent to glioblastoma cells, followed by RT-qPCR to evaluate effects on mRNA expression. Our analysis revealed decreased expression levels of self-renewal and pluripotency markers SOX2, NANOG and OCT in MTX treated T-ALL cells grown in co-culture compared to treated cells in suspension (Figure 4D). In contrast, NANOG was found to be upregulated in T-ALL cells in co-culture when not exposed to chemotherapy (Figure SF4A). These results suggest that CNS co-culture inhibits self-renewal of T-ALL cells during treatment, yet cells in co-culture show no loss in viability (Figure SF4D). Moreover, altered expression levels of cyclins and cyclin-dependent kinases (CDKs) involved in regulating cell cycle progression were identified in the co-culture system, with and without MTX exposure. Compared to culturing in suspension, co-culture of T-ALL cells both with and without MTX exposure, lead to inhibition of CDK2 and cyclin A2, which drives S/G2 transition, as well as CDK1, which together with Cyclin B is responsible for G2/M transition (Figure 4E, Figure SF4E) [24]. Furthermore, expression levels of Cyclin C and CDK3, essential for G0/G1 transition, were increased during CNS co-culture before treatment, with Cyclin C levels increasing also in co-culture during MTX exposure (Figure SF4E) [25]. Although MTX treatment efficiently reduced leukemia cell viability, cells grown in CNS co-culture showed an altered gene expression pattern compared cells in suspension, promoting cell survival. Together, this data indicates a shift towards cell cycle arrest and senescence of T-ALL cells when directly interacting with CNS derived cells in our glioma cell based co-culture system, potentially promoting therapy resistance.

## 4. DISCUSSION

Leukemia cells are capable of infiltrating and residing within the CNS, mainly entering via the blood leptomeningeal barrier or the blood cerebrospinal fluid barrier, where they interact with components of the microenvironment and remain sheltered from systemic treatment.

Leukemia cells can survive in the CNS niche, by hijacking the microenvironment and disrupting normal functions, promoting malignant transformation. These cells can then acquire quiescent properties, promoting a state of dormancy, and allowing survival in the new, unfavorable environment [26]. While the protective effects of the bone marrow niche have been widely studied, the mechanisms behind leukemia infiltration into the CNS, and the molecular changes that occur promoting leukemia cell survival in the CNS niche, remain unknown [23]. Due to the inability to accurately quantify CNS leukemic load and response to treatment, most CNS relapses occur in patients originally diagnosed as CNS negative [26]. In the light of this, no new drugs for CNS leukemia have been licensed since the 1960s [4]. Identifying specific molecular and epigenetic drivers to accurately measure CNS leukemic load could serve as solid biomarkers to predict CNS disease thus avoiding over or under treatment. Recent studies have linked interactions of leukemia cells and cells of the CNS niche, primarily meningeal cells, to increased survival and chemoresistance when cultured together *in vitro* [27]. Only when interrupting the interactions by applying adhesion inhibitors, the survival of leukemia cells in the co-culture decreased. Although research has shown that leukemia cells can resist treatment by interacting with components of the CNS niche, molecular aspects underlying these effects are yet to be defined [23].

We have identified a distinct differential gene expression pattern between pediatric T-ALL patients originally diagnosed as CNS3 at time of diagnosis, and CNS relapse, compared to CNS1 T-ALL patients. In samples with CNS involvement at time of diagnosis, an overall trend of gene downregulation was observed compared to patients with no CNS infiltration. However, in CNS relapse samples the opposite occurred, with an overall upregulation of differentially expressed genes. In CNS infiltrated T-ALL samples, RASSF2, a tumor suppressor with pro-apoptotic capacities often inactivated in cancer [11] and the proto-oncogene MDM2 were downregulated, indicating alterations of leukemia cell survival induced by the CNS niche. This was further typified by an upregulation of the long isoforms of the BCL2 family of apoptosis regulatory genes BCL2 and BCLX, which are known for their important role in promoting cell survival [28]. In line with this, there is a downregulation of the prop-apoptotic short isoforms in all three stages of CNS involved T-ALL, as well as CNS relapse.

Alterations in tumor suppressor genes, such as SETD2, have been associated with aggressive disease and relapse in leukemia patients [14]. Although the underlying mechanism of SETD2 remains largely unexplored, functional loss such as mutations in the SETD2 gene have been suggested to play a role in tumorigenesis, progression, and chemoresistance, thus leading to an unfavorable prognosis [20]. SETD2 has previously been recognized to as a tumor suppressor by interacting with p53 [14]. In line with our findings, decreased SETD2 expression is associated with increasing level of CNS involvement and is at its lowest range in CNS3 and CNS relapse. Interestingly, we have identified a significant negative correlation between SETD2 and TP53 in CNS relapsed patients. In contrary, a positive correlation between SETD2 and the E3 ubiquitin ligase MDM2 is seen. MDM2 carries out its function by binding to the N-terminal transactivation domain of p53 resulting in inhibition of transcriptional activation of p53 [29], thus promoting tumor formation. Our results are further potentiated mechanistically by overexpression of SETD2, which show direct impact on both MDM2 and TP53, as well as the apoptotic regulators BCL2 and BCLX. Additionally, our glioma co-culture system further supports a microenvironmental involvement in the malignant regulation of MDM2 and TP53, where an upregulation of MDM2 followed by a downregulation of TP53 in T-ALL cells directly interacting CNS-derived cells. Although the role of SETD2 and the H3K36me3 histone mark have been debated, a distinct upregulation of the histone mark is seen in the co-culture system. The co-culture system also highlights the importance of the microenvironmental niche with increased levels of adhesion markers such as CD44. CD44 have previous been described to play an important role in leukemia disease progression, and inhibition of CD44 in animal models have shown a release of the leukemia stem cells from the malignant niche, making them more accessible for standard chemotherapy [19].

Finally, we performed treatment with intrathecal prophylactic chemotherapy (MTX) in our co-culture system. A decrease in pluripotency markers as well as a halted cell cycle progression was observed, compared to MTX-treated cells grown outside the CNS co-culture. Members of the CDK family are impacted which are critical in the cell cycle regulation. Also, dysregulation of cyclins was identified, with downregulation of Cyclin A, which controls CDKs. However, Cyclin C, which plays a tumor suppressive role in T-ALL by controlling NOTCH1 levels, is upregulated [30]. Together, our glioma cell co-culture system led to reduced proliferation and self-renewal of MTX-treated T-ALL cells when in direct contact with CNS derived cells, causing cell cycle arrest and senescence, thus promoting survival and therapy resistance.

In conclusion, our findings indicate that the CNS niche induces gene expression changes in leukemia cells, leading to a differential gene expression pattern in CNS infiltrated T-ALL promoting cell survival, chemoresistance and disease progression. Moreover, interactions between leukemia cells and the CNS microenvironment induce epigenetic alterations leading to changes in gene regulation due to various chromatin and histone modifications. These changes in gene expression can be utilized to predict CNS infiltration, thus avoiding over or under treatment. Genetic drivers of disease progression could potentially serve as therapeutic targets, altering leukemia cells in the CNS niche to become more sensitive to current therapeutic strategies.

## ACKNOWLEDGEMENT

This work was supported by the Swedish Childhood Cancer Foundation (Barncancerfonden; F.H) and Märta och Gunnar V. Philipson Foundation (A.N, F.H).

The TARGET Initiative (RNA-sequencing data-set): The results published here are in whole or part based upon data generated by the Therapeutically Applicable Research to Generate Effective Treatments (TARGET) initiative, phs000218, managed by the NCI. The data used for this analysis are available at https://portal.gdc.cancer.gov/projects/TARGET-ALL-P2.

Information about TARGET can be found at http://ocg.cancer.gov/programs/target. Part of the data handling was enabled by resources at project number SNIC-2021/22-48 provided by the Swedish National Infrastructure for Computing (SNIC) partially funded by the Swedish Research Council through grant agreement no. 2018-05973.

## AUTHORSHIP STATEMENT

Sabina Enlund, Amanda Ramilo Amor, Indranil Sinha, Konstantinos Papadakis, Christina Neofytou and Frida Holm collected and analyzed the data. Sabina Enlund and Frida Holm conceptualized the project, analyzed data, and wrote the paper. Anna Nilsson, Ola Hermanson and Qingfei Jiang contributed with scientific expertise and were involved in writing and reviewing the paper.

## FIGURE LEGEND

**Supplemental figure 1**.

**A)** Venn diagram showing distribution of significantly expressed genes (p<0.05) in diagnosis samples divided by CNS status (without novel transcripts).

**B)** Percentage of significantly differentially expressed genes (left pie chart) and distribution of up and downregulated genes (right pie chart) in CNS3 diagnosis patients compared to CNS1 diagnosis patients.

**C)** Heat map of 50 differentially expressed genes in CNS3 versus CNS1 diagnosis patients, data displayed as log values collapsed to symbols.

**D)** Waterfall plot of showing top differentially expressed genes in CNS relapse versus CNS1 diagnosis patients, displayed as L2FC calculated from normalized FPKM values based on RNA-sequencing analysis (p<0.05).

**Supplemental figure 2**.

**A)** GSEA enrichment plot of cell adhesion molecules in CNS relapse versus diagnosis patients without CNS involvement, showing the profile running ES score and positions of gene set members on the ranked-ordered list. GSEA was performed using KEGG pathways.

**B)** GSEA enrichment plot of Notch signalling pathway in CNS relapse versus diagnosis patients without CNS involvement.

**C)** GSEA enrichment plot of Wnt signalling pathway in CNS relapse versus diagnosis patients without CNS involvement.

**D)** GSEA enrichment plot of apoptosis in CNS relapse versus diagnosis patients without CNS involvement. E) RNA-sequencing analysis of normalized FPKM values showing decreased expression of BCL2 in CNS relapsed T-ALL and no significant difference of BCLX expression levels. (Significance calculated by two-tailed, unpaired Student’s t-test. *p<0.05, **p<0.01).

**Supplemental figure 3**.

**A)** RNA-sequencing analysis of normalized FPKM values showing decreased expression of KAT6A in CNS involved T-ALL compared to non-CNS involved diagnosis patients.

**B)** RNA-sequencing correlation analysis of SETD2 and TP53 (left graph, r=0.072) and MDM2 (right graph, r=0.029) showing no significant correlation in diagnosis patients with no CNS involvement. (Correlation was calculated using Pearson correlation coefficients, two-tailed with 95% confidence interval).

**C)** RNA-sequencing analysis of normalized FPKM values showing decreased expression of EP300 (left graph) and SIRT1 (right graph) CNS involved T-ALL compared to non-CNS involved diagnosis patients. (Significance calculated by two-tailed, unpaired Student’s t-test. *p<0.05).

**D)** RNA-sequencing correlation analysis of SETD2 and EP300 (left graph) showing a positive correlation (r=0.817, ****p=<0.0001) and SIRT1 (right graph) showing a positive correlation (r=0.464, ***p=<0.001) in all included T-ALL patients. (Correlation was calculated using Pearson correlation coefficients, two-tailed with 95% confidence interval).

**E)** Schematic figure of SETD2 overexpression vector.

**F)** SETD2 mRNA expression levels (fold change versus empty vector control, EV=empty vector) 24 hours post SETD2 overexpression by transfection in MOLT16, Jurkat and CCL-119 cells. G) mRNA expression levels (fold change versus empty vector control) of BCL2 and BCLX short and long isoforms after SETD2 overexpression in MOLT16 cells.

**Supplemental figure 4**.

**A)** RT-qPCR analysis of T-ALL cell lines in CNS co-culture showing increased expression (fold change) of ICAM and NANOG, but not no difference of SETD2 compared to cells grown in suspension. (Significance calculated by two-tailed, unpaired Student’s t-test. *p<0.05, **p<0.01).

**B)** Chromatin Immunoprecipitation coupled with RT-qPCR showing increased expression of H3K36me3 on the BCLX promoter region in cell lines grown in CNS co-culture compared to cells grown in suspension. (Significance was calculated by two-way ANOVA, ***p<0.001).

**C)** RT-qPCR analysis of T-ALL cell lines in CNS co-culture showing increased expression (fold change) of BCLX compared to cells grown in suspension. (Significance calculated by two-tailed, unpaired Student’s t-test. **p<0.01).

**D)** Percentage of viable T-ALL cells (MOLT16, Jurkat, CCL-119 and SUP-T1) grown in co-culture and suspension after 48 hours of MTX treatment.

**E)** RT-qPCR analysis of T-ALL cell lines in CNS co-culture and suspension with and without 48 hours of 2uM MTX treatment, showing increased expression (relative to HPRT) of Cyclin C in non-treated and MTX treated T-ALL cell lines in grown co-culture and increased expression of CDK3 in T-ALL cell lines grown in co-culture, compared to cells grown in suspension, as well as decreased expression of cyclin B1 in MTX treated T-ALL cell lines grown in co-culture and in suspension compared to non-treated cells. (Significance calculated by two-tailed, unpaired Student’s t-test. *p<0.05, **p<0.01, ***p<0.001).

