## Supplementary figures and images for "The CNS Microenvironment Promotes Leukemia Cell Survival by Disrupting Tumor Suppression and Cell Cycle Regulation in Pediatric T-cell Acute Lymphoblastic Leukemia"

### Supplemental Figure 1

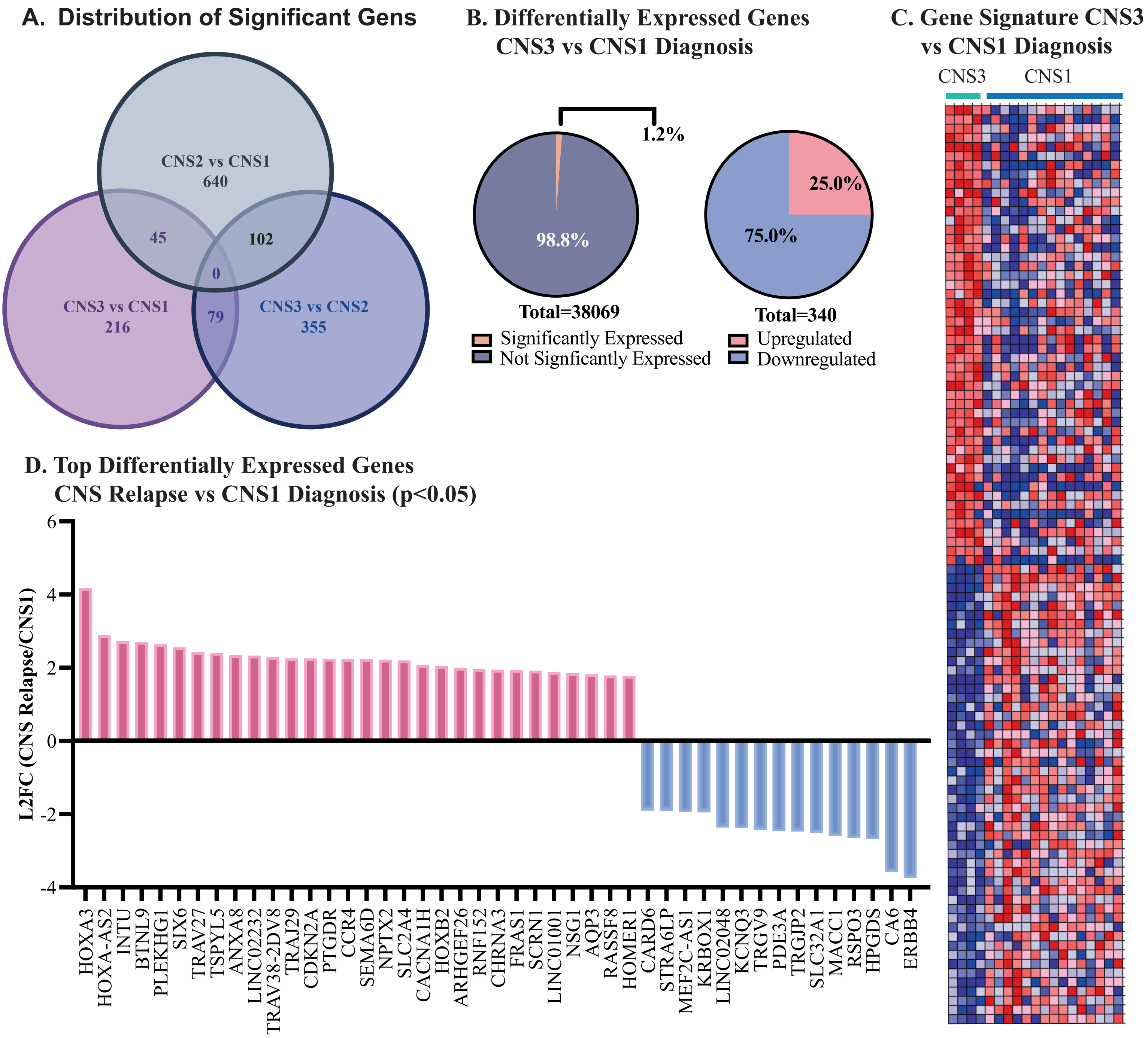

### Supplemental Figure 2

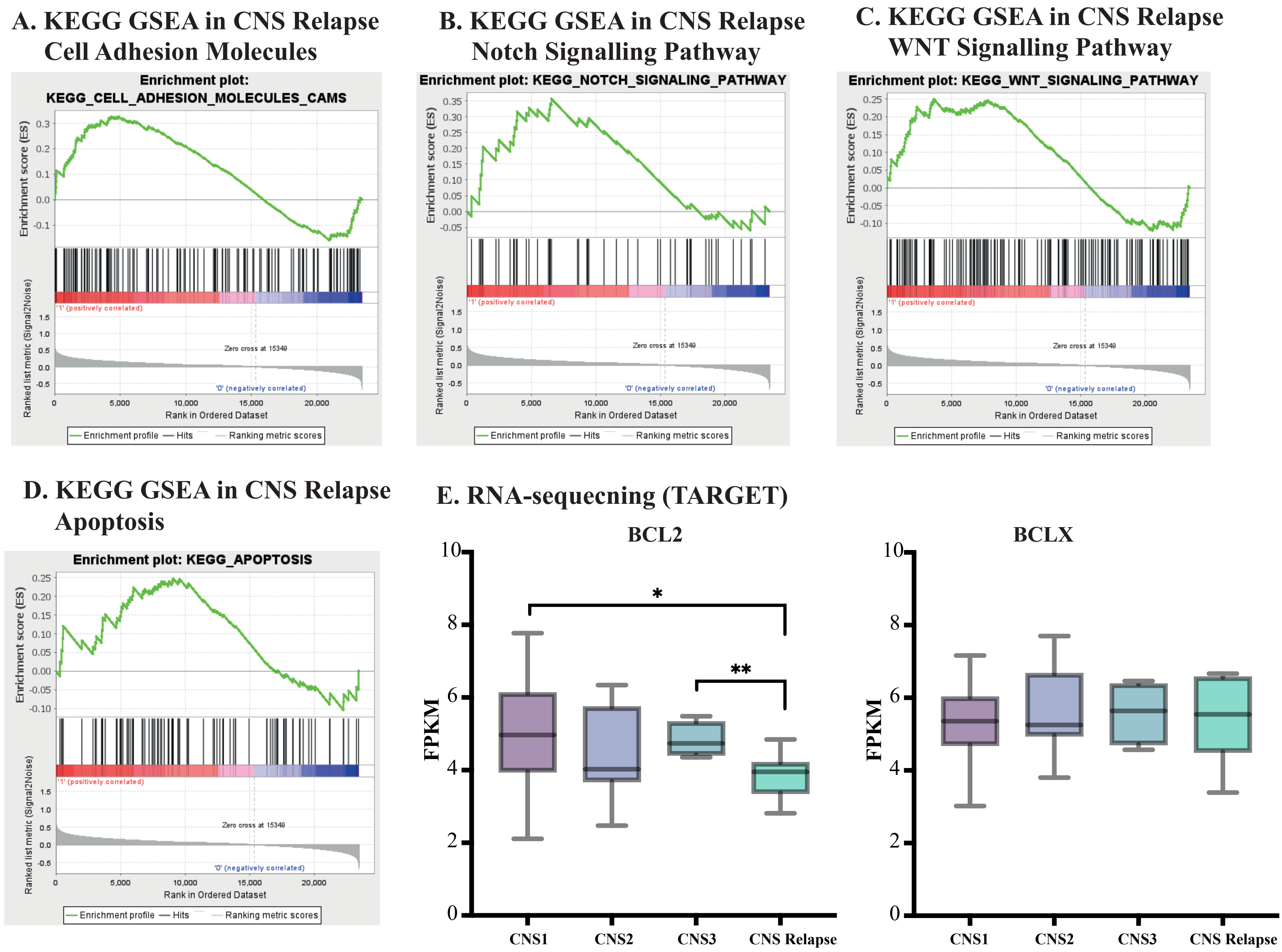

### Supplemental Figure 3

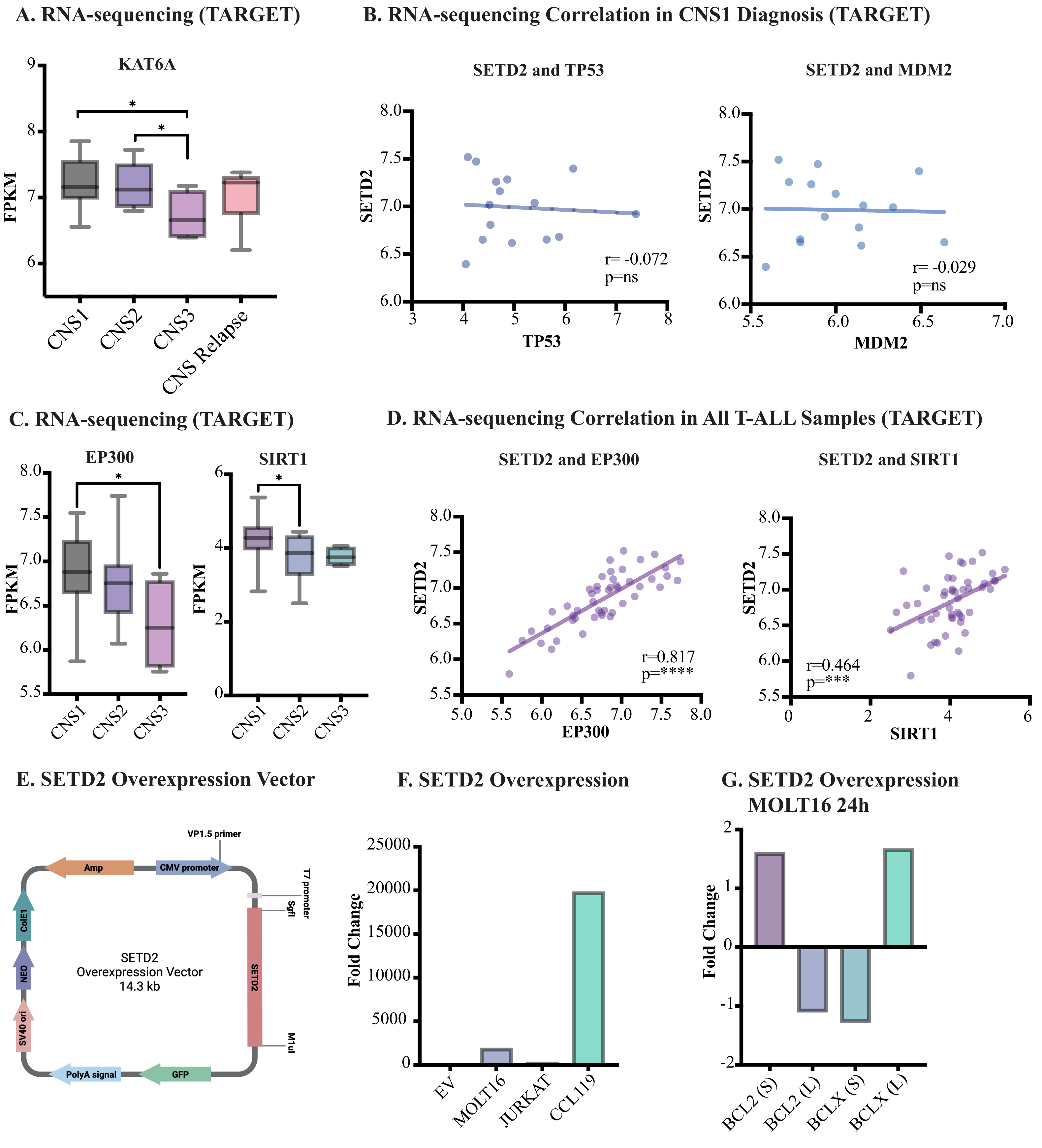

### Supplemental Figure 4

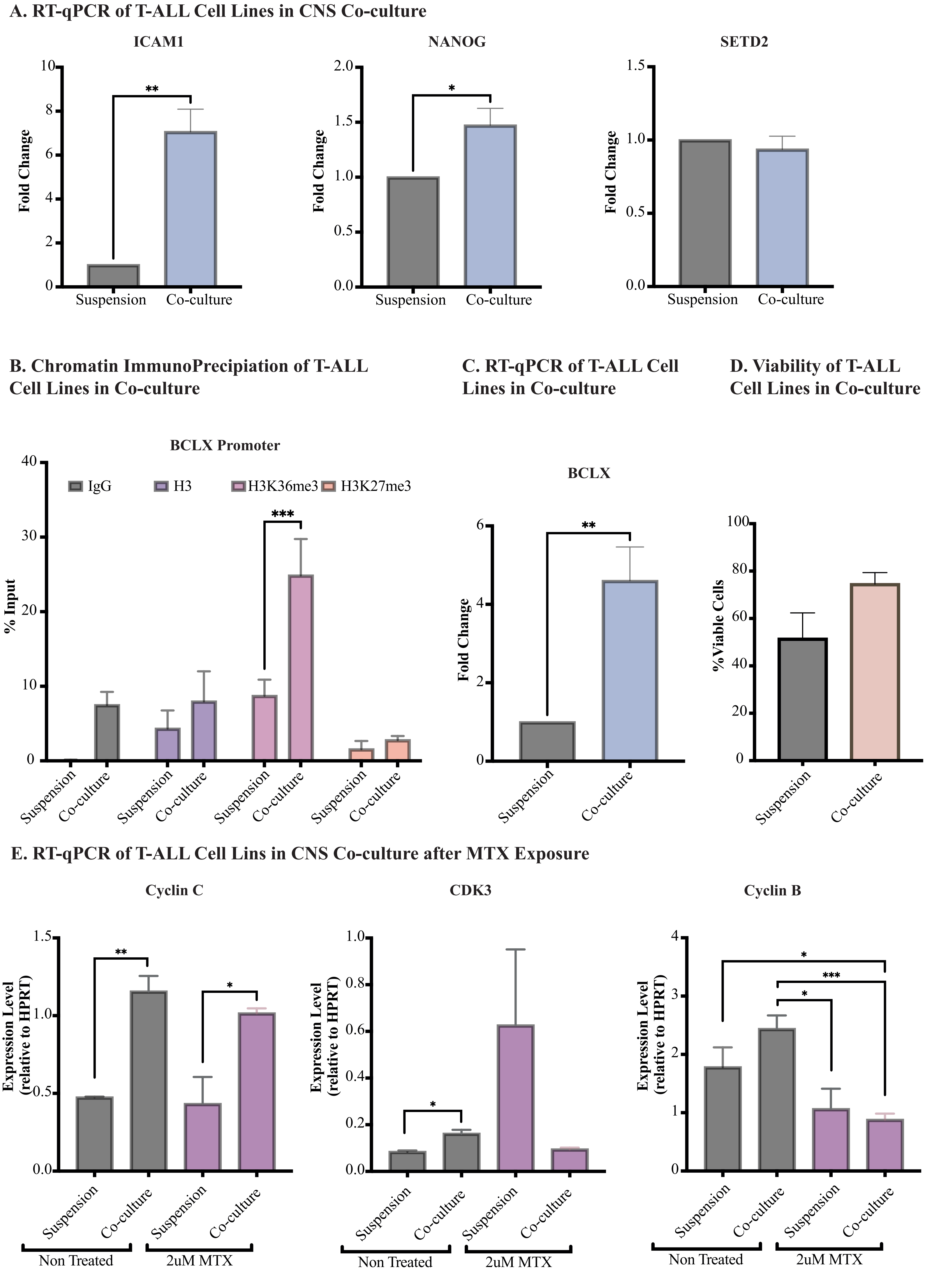
